# Resistance levels to the cassava green mite, *Mononychellus tanajoa*, in cassava germplasm (*Manihot esculenta*)

**DOI:** 10.1101/2020.08.26.268227

**Authors:** Jaime Marín, Arturo Carabali, James Montoya Lerma

## Abstract

The cassava green mite (CGM), *Mononychellus tanajoa* (Acari: Tetranychidae), is one of the main pests of cassava, causing direct damage by sucking the plant’s sap. Although the mite has a wide distribution in Latin America and Africa and a high potential to expand to Asia, limited information is available on *M. tanajoa* biology and life history parameters on its primary host. In this study, we quantified the levels of resistance of 10 cassava genotypes (i.e., NAT-31, ALT-12, ALT-6, COL-1505, ECU-72, ECU-160, PER-182, PER-335, 60444, CMC-40) based on the mite’s oviposition preference and development time in no-choice and choice bioassays. The genotype NAT-31 significantly differed from other genotypes for *M. tanajoa* development time and oviposition rate: each stage of the CGM life cycle appeared to be delayed in NAT-31, suggesting that NAT-31 resistance is mediated through a general reduction of CGM fitness on this genotype. Resistance in the remaining genotypes was variable in comparison to a susceptible (control) genotype. ECU-72, a parental line of NAT-31, present a difference related to oviposition preference, development time and sex ratio. These parameters allow the identification of different levels of resistance (antixenotic and antibiosis) when compared to the susceptible genotype. CGM displayed significantly different oviposition preference from the susceptible genotypes. Identification and characterization of resistance to CGM in cassava germplasm might be key to further advance knowledge about natural resistance mechanisms and develop strategies to introgress resistance to CGM in farmer- and industry-preferred cassava varieties.

## Introduction

Cassava (*Manihot esculenta* Crantz, Euphorbiaceae) is a woody perennial shrub originating from South America (Olsen and Schaal, 1999). With a total world production of over 280 million tons in 2012, it constitutes an essential source of carbohydrates for about 800 million people in tropical countries (FAO, 2012; FAOSTAT, 2009; Lebot, 2009). In addition to its use in human diets, cassava is widely used as a raw material in processed products as well as for the animal feed, ethanol and starch industries (Anggraini *et al*., 2009; Balagopalan 2002).

Approximately 200 species of arthropod pests are associated with cassava (Bellotti *et al*., 2002, 2010, 2012). Green mites [cassava green mite (CGM), *Mononychellus tanajoa* (Bondar), and *Mononychellus caribbeanae* (McGregor)], whiteflies (*Aleurotrachelus socialis, Bemisia tabaci* and *Aleurothrixus aepim*) and mealybugs (*Phenacoccus manihoti*) are among the most important arthropod pests infecting cassava (Parsa *et al*., 2015; Bellotti, 2002). These pest species, almost all native from the Neotropics, have adapted in various forms to the physical and biochemical defences of the plant, including its leaf pubescence and its laticiferous and cyanogenic compounds (Bellotti and Riis, 1994). Green mites and mealybugs negatively impact cassava yield by feeding on the terminal parts of the plants triggering cell death and reduced photosynthesis (Gomez *et al*., 2001). Field research has indicated that extended attacks (i.e., between 3 and 6 months) of *M. tanajoa* can cause up to 80% losses in root yields (Bellotti *et al*., 2012). *Mononychellus tanajoa* attacks are favoured by dry conditions, in unfertil plants, inadequate fertilisation and the presence of weeds (Bellotti *et al*., 2012). Whereas *M. tanajoa* presently affects several of the world’s prime cassava-growing areas, climate change and continued global spread have been predicted to further exacerbate mite pest problems in the African rift valley, the Mato Grosso in Brazil, northern South America and Southeast Asia (Herrera *et al*., 2011). The vast majority of cassava farmers tend to use insecticides to control CGM (CIAT, 2006; Arias, 1995). However, efficacy of spraying against this mite is usually limited, and multiple applications are required to keep CGM populations under control, making the crop economically unsustainable in regions where green mites are endemic (Panda and Khush, 1995). Biological control relying on the use of naturally occurring predators and entomopathogens as well as the deployment of mite-resistant cassava varieties are effective and complementary methods that may be used to manage *M. tanajoa* (Bellotti *et al*., 2012).

Wild relatives of cassava constitute important sources of genes for resistance against cassava arthropod pests (Vargas *et al*., 2002; Burbano *et al*., 2007; Carabali *et al*., 2010a, b; Parsa *et al*., 2015). Moderate to high levels of resistance to green mites, whiteflies and mealybugs were identified in inter-specific hybrids of *M. esculenta* subsp. (CIAT, 2006; Carabalí *et al*. 2010a, 2013). Furthermore, resistance to CGM was transferred to F_1_ inter-specific hybrids, suggesting a simple inheritance of this trait (A. Bellotti and M. Fregene, pers. comm.). However, the long reproductive cycle and lengthy time required to develop new cassava varieties (8-10 years) often discourages the use of wild species in conventional cassava breeding programs (Rudy *et al*., 2010; Legg *et al*., 2006). A preliminary assessment by Parsa *et al*. (2015) reported 33 potential sources of resistance to *M. tanajoa* in the cassava germplasm. Robust assessment of resistance levels to CGM in the cassava germplasm as well as identification of different types of resistance are urgently needed to develop and deploy sustainable management of CGM resistance in the field. Characterization of the CGM resistance is also essential to help understanding the associated physiological and phenotypic traits. In the present study we report on the assessment of resistance levels to CGM of selected cassava breeding lines. We used bioassays and measured biological parameters such as oviposition preference, development time and sex ratio to estimate the resistance level and to characterize the impact of resistant cassava host on CGM.

## MATERIALS AND METHODS

### Plants and mites

The study was conducted in 2012 in glasshouse facilities at the Universidad del Valle (Univalle) in Cali, Colombia (28 ± 2°C, 70 ± 5% relative humidity [RH]). Ten genotypes of *M. esculenta* (i.e., NAT-31, ALT-12, ALT-6, COL-1505, ECU-72, ECU-160, PER-182, PER-335, 60444, and CMC-40) were obtained from the CIAT germplasm bank. Genotypes were selected based on their potential or known levels of arthropod resistance/susceptibility (Table 1). CMC-40 accession was used as a susceptible control and was also used as host plant for CGM colony maintenance (Bellotti, 2002, Bellotti *et al*., 2010, 2012). All the genotypes were established *in vitro* and then planted in sterile soil in plastic pots and kept in a glasshouse at 30 ± 2°C and 70 ± 5% RH. Plants were irrigated 3× per week and the plants did not receive pesticide or fertilizer applications. Six-month-old plants were used for the bioassays. A stock colony of *M. tanajoa*, isolated in the cassava fields at CIAT (Cali, Colombia), was established and reared on CMC-40 plants under controlled conditions (28 ± 2°C, 70 ± 5% RH and L12:D12 photoperiod) for the bioassays. Cassava green mites were placed on fresh cassava plants every 20 days. Hybrid vigor was guaranteed bringing mites from the field every 2 months.

**Table 1.** Recorded cassava genotypes with variable levels of resistance or tolerance to arthropod pests.

| Accession | Mites | Whiteflies | Other pests | Resistance/tolerance | References |
| --- | --- | --- | --- | --- | --- |
| ALT-6 | *Mononychellus tanajoa* |  |  | Tolerance | CIAT, 2004 |
| ALT-12 | *M. tanajoa* |  |  | Tolerance | CIAT, 2004 |
| COL-1505 | - | - | - | - | - |
| ECU-72 |  | *Aleurotrachelus socialis*, *Bemisia tabaci* |  | Resistance | Bellotti and Arias, 2001; Gómez, 2004; Carabalí *et al.*, 2009; Bohorquez, 2009; Omongo *et al.*, 2012; Parsa *et al.*, 2015 |
| ECU-160 | *M. tanajoa* |  |  | Tolerance | Burbano *et al.*, 2007 |
| NAT-31 (CG 489-31) |  | *A. socialis* |  | Resistance | Vargas *et al.*, 2002 |
| PER-182 | *M. tanajoa* |  |  | Tolerance | Boaventura *et al.*, 2006 |
| PER-335 | *M. tanajoa* | *A. socialis* |  | Tolerance | Bellotti and Arias, 1981; Boaventura *et al.*, 2006 |
| 60444 |  |  | *Erinnyis ello* | Tolerance | Herrera *et al.*, 2011; CIAT, 2004 |

### Bioassays

Mite performance on the selected genotypes was evaluated using the following biological parameters: host plant selection, oviposition preference, development time and offspring sex ratio. First, in a preliminary trial, the 10 genotypes were screened, followed by a second trial using the five genotypes that were tolerant to CGM attack in the first trial. Due to reduced establishment rate using stem propagation for ALT-6, this genotype was not included in the second screening assay. In all cases, two sets of bioassays (choice and no-choice) were conducted.

Oviposition preference assays were based on choice and no-choice tests. The choice assay consisted of facing susceptible CMC-40 (control) genotype with the other nine genotypes in a Petri dish (200 × 15 mm). Lobes of each genotype were placed on top in a circle on foam moistened with water. In the choice assay, the position of the materials in the Petri dish was rotated clockwise, in order to guarantee independence and their random distribution. In the no-choice assays, 10 lobes of the same genotype were placed in a Petri dish. Experimental unit was the Petri dish with 10 lobes in the first assay (in total five Petri dishes, 50 replicates). For the second assay, the experimental unit was the same Petri dish but with five lobes (in total 10 Petri dishes, 50 replicates).

### Oviposition preference

For the choice assays, a leaf lobe (average length 3 cm) of each genotype was placed on humid foam and then arranged in a circle inside a Petri dish (200 × 15 mm) as shown in Figure 1. Twenty pairs (male and female) of CGM adults, 24 h post emergence, were placed on CMC-40 leaf lobes at the centre of a Petri dish (50 for each genotype, 10 petri dishes with 10 lobes in each of the petri dishes.) (Figure 1A). For the no-choice assays, a similar procedure was followed with 20 pairs of mites per arena placed on CMC-40 leaf discs, but now each Petri dish contained only lobes of a single genotype (Figure 1B) (one experimental run in which 20 mite pairs were released within a single Petri dish).

**Figure 1.**
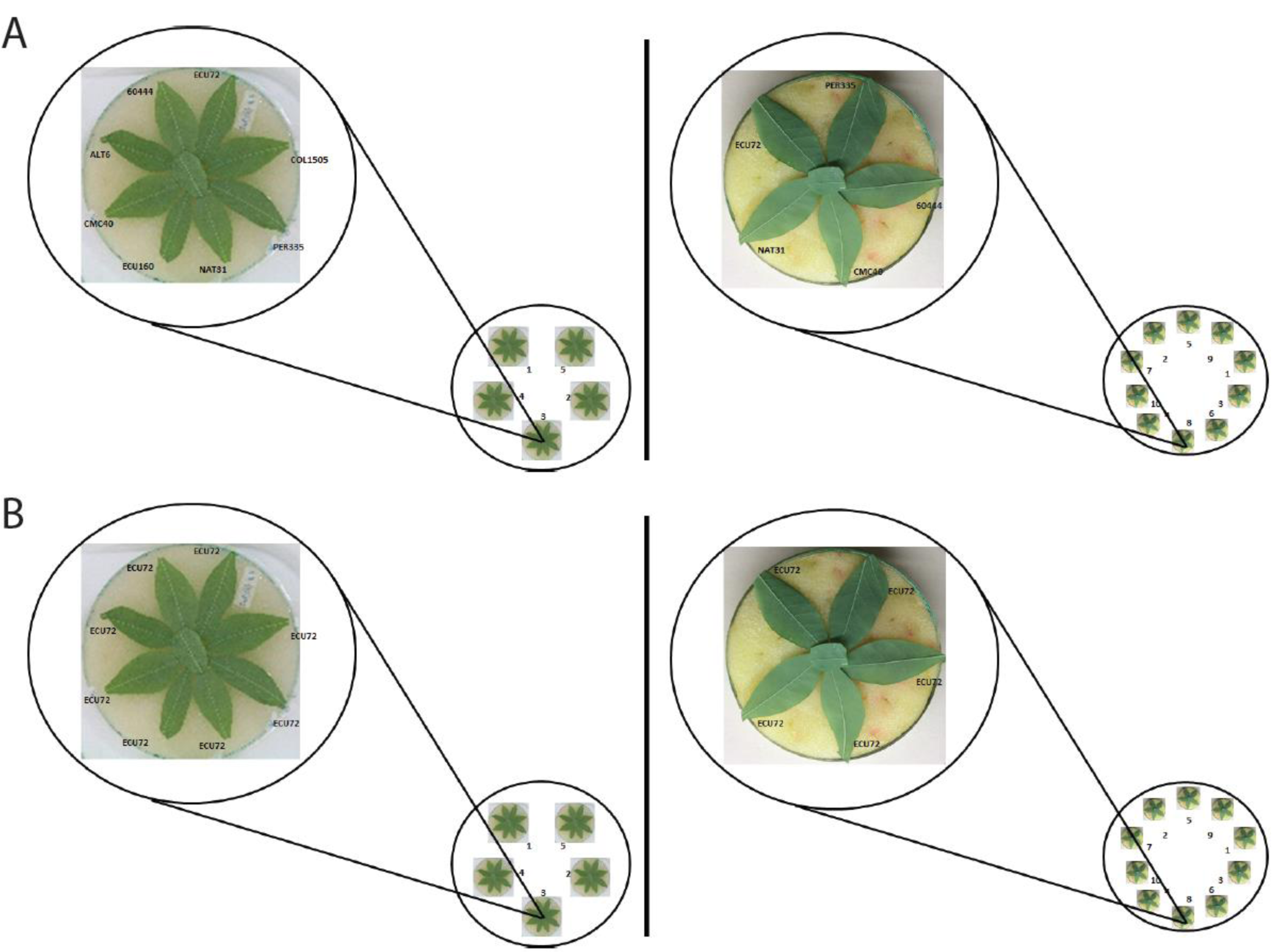
Design of the (A) choice and (B) no-choice assays for oviposition preference of cassava green mite on leaf lobes of various cassava genotypes. Initially mite responses were screened on leaves of 10 genotypes (left) and subsequently responses were assessed on five genotypes (right) after 1, 2 and 8 days.

Oviposition was evaluated for eight (8) days and estimated as the average number of eggs laid per female on 50 lobes for each genotype (choice assays) and the average number of eggs laid per female on each genotype (no-choice). In each case, this was asseessed counting the total number of eggs laid per female during twice observations per day. The two observations did not show statistical differences; hence for the analysis the total diaries were used. The free and non-free preference tests were carried out for 10 and 12 days. Count started from the petiole (upper rib) and continued in the lower part of the rib. The number of eggs at 24h and 48h was compared in order to detect any early difference among genotypes.

### Development time and sex ratio

Male and female mite adults, obtained from 6-month-old CMC-40 plants, were transferred to the underside of leaf lobes obtained from 6-month-old plants, kept on humid foam inside Petri dishes (200 × 15 mm). The average size of the lobes was 3 cm. After 12 h, the adults were eliminated, and eggs were randomly selected and removed with a needle and a fine brush. Only one egg was left to continue its development until reach adulthood. Observations were done every 2 h recording its physiological stage. Fifty lobes of each genotype were evaluated for egg to adult development time and offspring sex ratio.

### Statistical analysis

Differences among the mean values in the no-choice and choice tests on the various genotypes were analysed using one-way ANOVA, followed by Tukey test for multiple mean comparison tests. ANOVA was used to detect differences in fecundity, development time and sex ratios. All analyses used the R statistical program (R Development Core Team, 2014). The level of significance was 5%. All biological parameters evaluated had 50 technical replicates in all assays.

## RESULTS

### Oviposition preference

Significant differences were recorded between the numbers of eggs on the 10 genotypes, as compared with CMC-40 (ANOVA P <0.0001, followed by Tukey P <0.05) (Table 2). The oviposition rate ranged from 1.5±0.09 to 27.1±1.98 eggs/8 days for the assessed genotypes (Table 2). Female mites showed the highest oviposition rates on genotype 60444 (27.1± 1.98 eggs) with average number of eggs approximately twice compared to the susceptible check CMC-40 (14.8± 1.02 eggs). NAT-31 and ALT-6 displayed low oviposition rates (1.5± 0.09 eggs and 3.3±0.16 eggs, respectively, Table 2). Significant differences were recorded among genotypes (*P* < 0.05) at 24h and 48h (Table 2). A group of genotypes (PER-182, 60444, CMC-40) are preferred at 24h while presence of females was substantially delayed on NAT-31, ALT-6, ECU-72, ECU-160.

**Table 2.** Choice assay for oviposition preference of *Mononychellus tanajoa*: mean (± SE) number of eggs laid after 24 and 48 h and preference for oviposition measured by no. eggs/genotype and average oviposition rate (N=500).

| Accession | First trial |  |  |  | Second trial |  |
| --- | --- | --- | --- | --- | --- | --- |
|  | No. eggs after 24 h | No. eggs after 48 h | No. eggs after 8 days | Oviposition rate (no. eggs/ Per day and per females) | No. eggs after 8 days | Oviposition rate (no. eggs/ Per day and per females) |
| 60444 | 6.14±7.93<sup>a</sup> | 8.88±11.93<sup>a</sup> | 27.1±1.98<sup>a</sup> | 9.9±0.27<sup>a</sup> | 31.11±1.20<sup>a</sup> | 19.12±3.25<sup>a</sup> |
| PER-335 | 2.66±4.32<sup>bc</sup> | 5.7±9.0<sup>b</sup> | 17.5±1.39<sup>ab</sup> | 12.6±0.3<sup>ab</sup> | 29.72±1.04<sup>a</sup> | 20.16±3.67<sup>a</sup> |
| PER-182 | 1.6±3.2<sup>cd</sup> | 3.54±6.62<sup>bc</sup> | 11.0±0.81<sup>bcd</sup> | 2.9±0.19<sup>bcd</sup> | - | - |
| ALT-12 | 1.44±2.43<sup>cd</sup> | 4.16±5.57<sup>bc</sup> | 13.7±1.01<sup>b</sup> | 2.74±0.27<sup>b</sup> | - | - |
| ALT-6 | 1.8±3.07<sup>cd</sup> | 1.86±3.05<sup>cd</sup> | 3.3±0.16<sup>cd</sup> | 0.8±0.005<sup>cd</sup> | - | - |
| COL1505 | 2.84±5.6<sup>bc</sup> | 6.32±9.86<sup>ab</sup> | 12.4±0.79<sup>bc</sup> | 2.8±0.34<sup>bc</sup> | - | - |
| ECU-160 | 1.46±4.15<sup>cd</sup> | 2.32±4.6<sup>cd</sup> | 9.2±0.81<sup>bcd</sup> | 2.3±0.08<sup>bcd</sup> | - | - |
| NAT-31 | 0.28±0.9<sup>d</sup> | 0.62±1.74<sup>d</sup> | 1.5±0.09<sup>d</sup> | 0.42±0.03<sup>d</sup> | 7.90±0.29<sup>b</sup> | 5.34±0.99<sup>b</sup> |
| ECU-72 | 1.54±2.89<sup>cd</sup> | 4.7±7.97<sup>bc</sup> | 12.9±0.93<sup>bc</sup> | 3.2±0.31<sup>bc</sup> | 35.42±1.21<sup>a</sup> | 23.70±4.3<sup>a</sup> |
| CMC-40 | 4.0±5.8<sup>b</sup> | 5.46±6.1<sup>b</sup> | 14.8±1.02<sup>b</sup> | 3.7±0.14<sup>b</sup> | 29.12±1.40<sup>a</sup> | 14.92±2.26<sup>a</sup> |
| F | 6.948 | 4.6985 | 10.851 |  | 7.7825 |  |
| d.f. | 9 - 490 | 9 - 490 | 9 - 490 |  | 4 - 245 |  |
| P | <1.855e<sup>-09</sup> *** | <5.331e<sup>-06</sup> *** | 1.889e<sup>-15</sup> *** |  | 6.412e<sup>-06</sup> *** |  |

Means within a column followed by the same letter are not significantly different (Tukey's HSD test: P>0.05).

Ten genotypes with contrasting oviposition rates were selected and tested with choice and no choice bioassays. The experiment confirmed the second trial, oviposition rates ranged between 5.34± 0.99 and 23.7± 4.3 eggs/female/8 days for choice bioassays and 4.4±0.28 and 13.34±2.97 eggs/female/8days for no choice bioassays (Tables 2 and 3). Oviposition rates (number of eggs/genotype) on NAT-31 and ALT-6 had, respectively, 79.2% and 82% reduction when compared to the oviposition rate of the susceptible check CMC-40 (Table 2). Significant differences were noted between the numbers of eggs on the four genotypes compared with those on CMC-40 (*P* < 0.05) (Figure 3). Oviposition ranged from 7.9±0.29-35.42±1.21 eggs/8 days in the second choice bioassays and 4.67±0.49-14.03±0.97 eggs/8 days in the no choice bioassay (Tables 2 and 3).

**Table 3.**
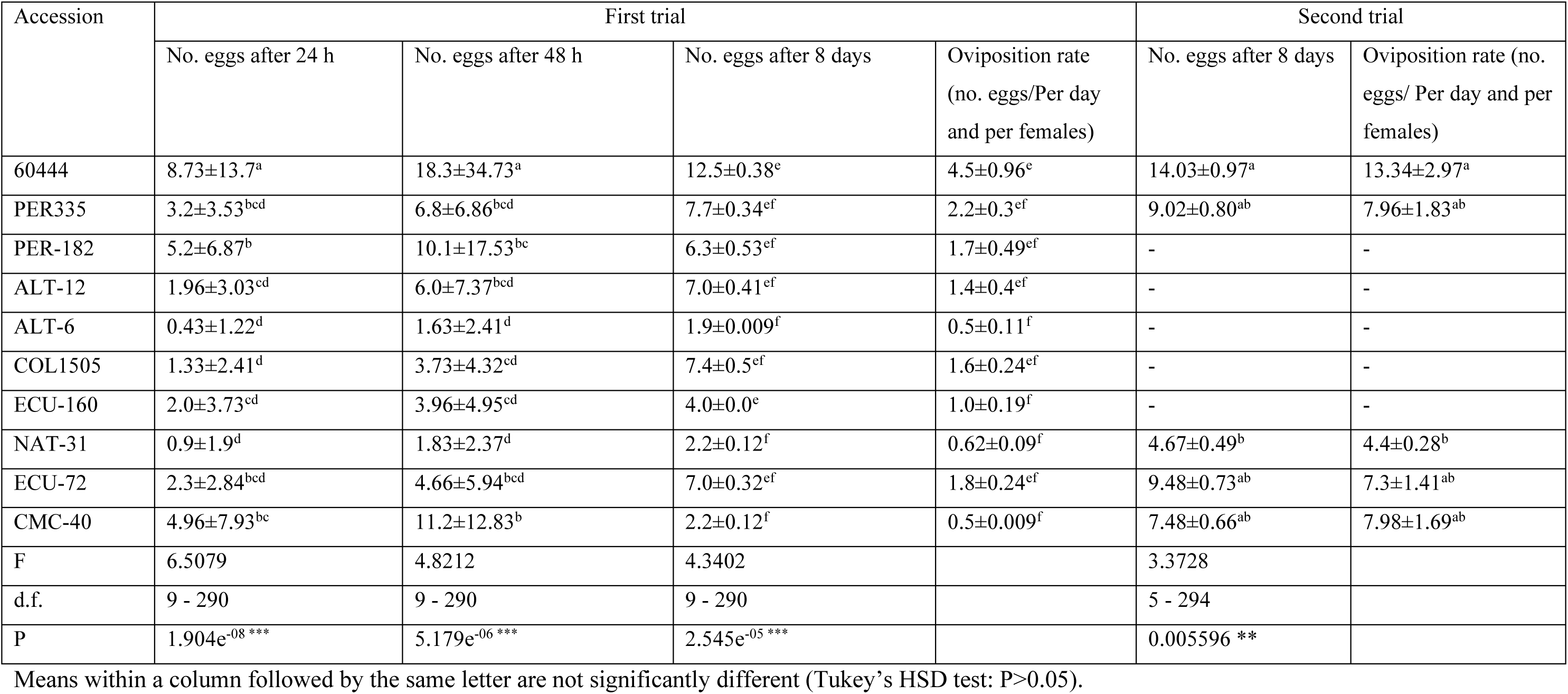
No-choice assays for oviposition preference of *Mononychellus tanajoa*: mean (± SE) number of eggs laid after 24 and 48 h and preference for oviposition measured by No. eggs/genotpype and average oviposition rate of *M. tanajoa* a no-free choice (N=500).

| Accession | First trial |  |  |  | Second trial |  |
| --- | --- | --- | --- | --- | --- | --- |
|  | No. eggs after 24 h | No. eggs after 48 h | No. eggs after 8 days | Oviposition rate<br>(no. eggs/Per day<br>and per females) | No. eggs after 8 days | Oviposition rate (no.<br>eggs/ Per day and per<br>females) |
| 60444 | 8.73±13.7<sup>a</sup> | 18.3±34.73<sup>a</sup> | 12.5±0.38<sup>e</sup> | 4.5±0.96<sup>e</sup> | 14.03±0.97<sup>a</sup> | 13.34±2.97<sup>a</sup> |
| PER335 | 3.2±3.53<sup>bcd</sup> | 6.8±6.86<sup>bcd</sup> | 7.7±0.34<sup>ef</sup> | 2.2±0.3<sup>ef</sup> | 9.02±0.80<sup>ab</sup> | 7.96±1.83<sup>ab</sup> |
| PER-182 | 5.2±6.87<sup>b</sup> | 10.1±17.53<sup>bc</sup> | 6.3±0.53<sup>ef</sup> | 1.7±0.49<sup>ef</sup> | - | - |
| ALT-12 | 1.96±3.03<sup>cd</sup> | 6.0±7.37<sup>bcd</sup> | 7.0±0.41<sup>ef</sup> | 1.4±0.4<sup>ef</sup> | - | - |
| ALT-6 | 0.43±1.22<sup>d</sup> | 1.63±2.41<sup>d</sup> | 1.9±0.009<sup>f</sup> | 0.5±0.11<sup>f</sup> | - | - |
| COL1505 | 1.33±2.41<sup>d</sup> | 3.73±4.32<sup>cd</sup> | 7.4±0.5<sup>ef</sup> | 1.6±0.24<sup>ef</sup> | - | - |
| ECU-160 | 2.0±3.73<sup>cd</sup> | 3.96±4.95<sup>cd</sup> | 4.0±0.0<sup>e</sup> | 1.0±0.19<sup>f</sup> | - | - |
| NAT-31 | 0.9±1.9<sup>d</sup> | 1.83±2.37<sup>d</sup> | 2.2±0.12<sup>f</sup> | 0.62±0.09<sup>f</sup> | 4.67±0.49<sup>b</sup> | 4.4±0.28<sup>b</sup> |
| ECU-72 | 2.3±2.84<sup>bcd</sup> | 4.66±5.94<sup>bcd</sup> | 7.0±0.32<sup>ef</sup> | 1.8±0.24<sup>ef</sup> | 9.48±0.73<sup>ab</sup> | 7.3±1.41<sup>ab</sup> |
| CMC-40 | 4.96±7.93<sup>bc</sup> | 11.2±12.83<sup>b</sup> | 2.2±0.12<sup>f</sup> | 0.5±0.009<sup>f</sup> | 7.48±0.66<sup>ab</sup> | 7.98±1.69<sup>ab</sup> |
| F | 6.5079 | 4.8212 | 4.3402 |  | 3.3728 |  |
| d.f. | 9 - 290 | 9 - 290 | 9 - 290 |  | 5 - 294 |  |
| P | 1.904e<sup>-08</sup> *** | 5.179e<sup>-06</sup> *** | 2.545e<sup>-05</sup> *** |  | 0.005596 ** |  |

Means within a column followed by the same letter are not significantly different (Tukey's HSD test: P>0.05).

**Figure 2.**
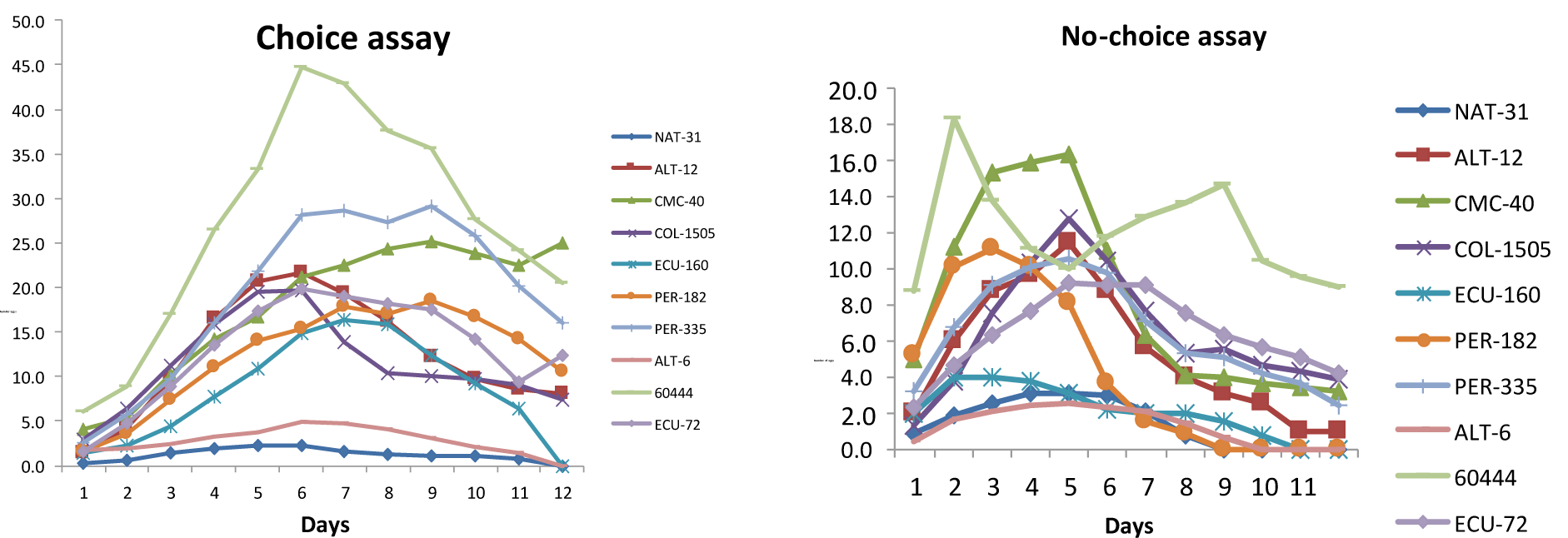
Test to free (A) and not free (B) preference for oviposition in 10 cassava genotypes, first selection. Evaluating in days the preference of the green mite to lay eggs.

**Figure 3.**
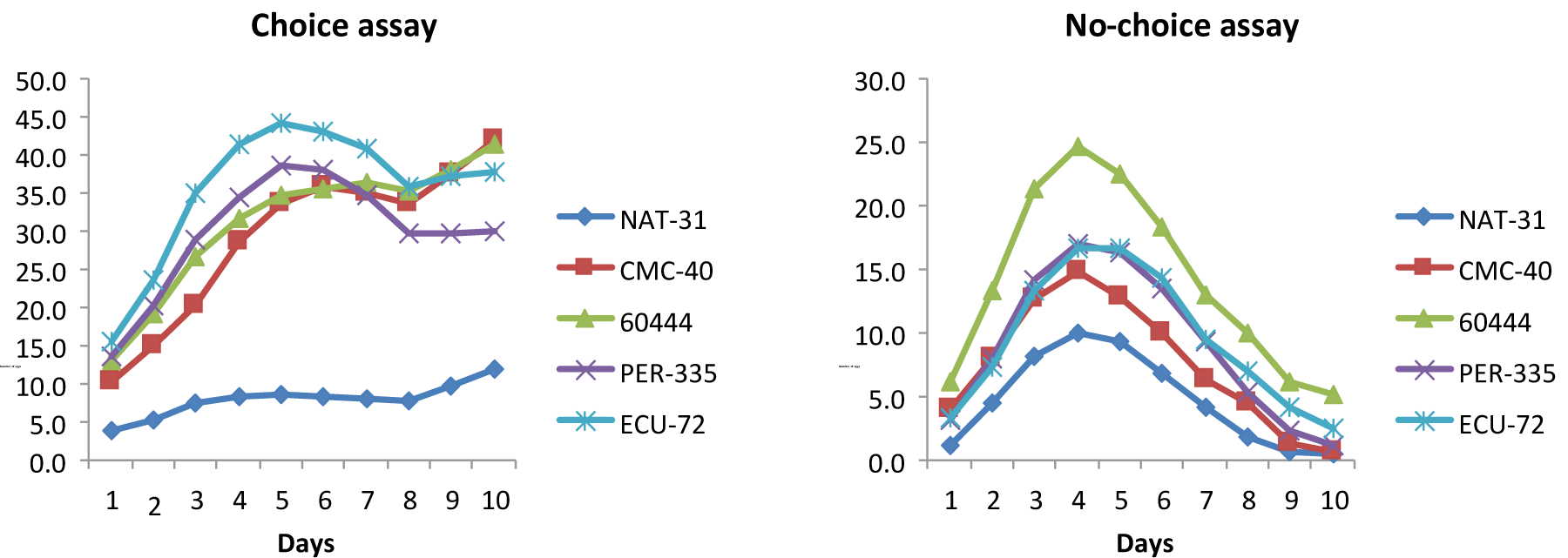
Test to free (A) and not free (B) preference for oviposition in 5 cassava genotypes, second selection. Evaluating in days the preference of the green mite to lay eggs.

In the second bioassay, the oviposition preference for genotypes 60444 (31.11± 1.20 eggs) and ECU-72 (35.42± 1.21 eggs) was similar to CMC-40 (29.12± 1.40 eggs) (Tables 2 and 3), and NAT-31 remained the least preferred (7.90± 0.29 eggs) (Table 2). There were significant differences in the number of eggs laid on NAT-31 and ALT 6 compared with the susceptible genotype (CMC-40) (Tables 2 and 3) (*P* < 0.05) used as control. The least preferred genotypes in the preliminary trial were NAT-31 (2.2±0.012 eggs) and ALT-6 (1.9±0.009 eggs), with 79.2% and 82% reduction in oviposition, respectively (Table 2). In the second trial, genotypes 60444 (14.03±0.97 eggs) and ECU-72 (9.48±0.73 eggs) had higher oviposition rates compared with CMC-40 (7.48±0.66 eggs) (Table 2). As in the preliminary selection, NAT-31 was the green mite least preferred accession (4.67±0.49 eggs). As in the first trial, at no choice assays, female mites preferred oviposit to at 24h and 48h but there were significant differences among genotypes (*P* < 0.05) (Table 2).

### Developmental time and sex ratio

Development time on the selected genotypes were similar with the exception of ECU-160, on which *M. tanajoa* had a significantly shorter development time as compared to the other genotypes (Table 4). Despite a relatively homogenous egg-to-adult development time in most genotypes, there were significant differences between genotypes for development time of particular stages. The CMC40 genotype in all experiments and evaluated parameters behaves as a slightly susceptible material when compared to 60444 (Tables 2 and 3; Figures 2 and 3). For this reason, material 60444 was selected as a susceptible material when dealing with pests such as the green mite. The CMC40 material can be compared to 60444, PER335 and ALT12. The genotypes NAT-31, ECU-72 y ALT-12 displayed either reduction or increase in life parameters of the acari. For instance, the developmental times were reduced when exposed to NAT-31 being 1.10±0.15 for protonymph and 0.67±0.09 for deutonymph but increased 6.22±0.30 for eggs; ECU-72 reduced 0.0±0.0 the teleiochrysalid stage while increased the larvae (1.55±0.14) and deutonymph (2.27±0.20). Finally, ALT-12 reduced in the egg stage (3.5±0.18) and increased for teleiochrysalid (1.05±0.06) and adults (3.50±0.19). The egg stage on the susceptible check CMC-40 was significantly shorter as compared to resistant genotypes. The short egg stage on CMC-40 was outbalanced by a longer protochrysalide stage (1.85±0.16 d) when compared, for example, to NAT-31 (0.7±0.07 d). On ECU-72, a genotype previously reported to be resistant genotype to pests (Bohórquez, 2009), the egg stage time (3.65±0.13 d) was comparable to the susceptible check CMC-40. The proportion of females (0.9:0.1) was not affected on two of the genotypes; ALT12 and COL1505 (Table 4).

**Table 4.**
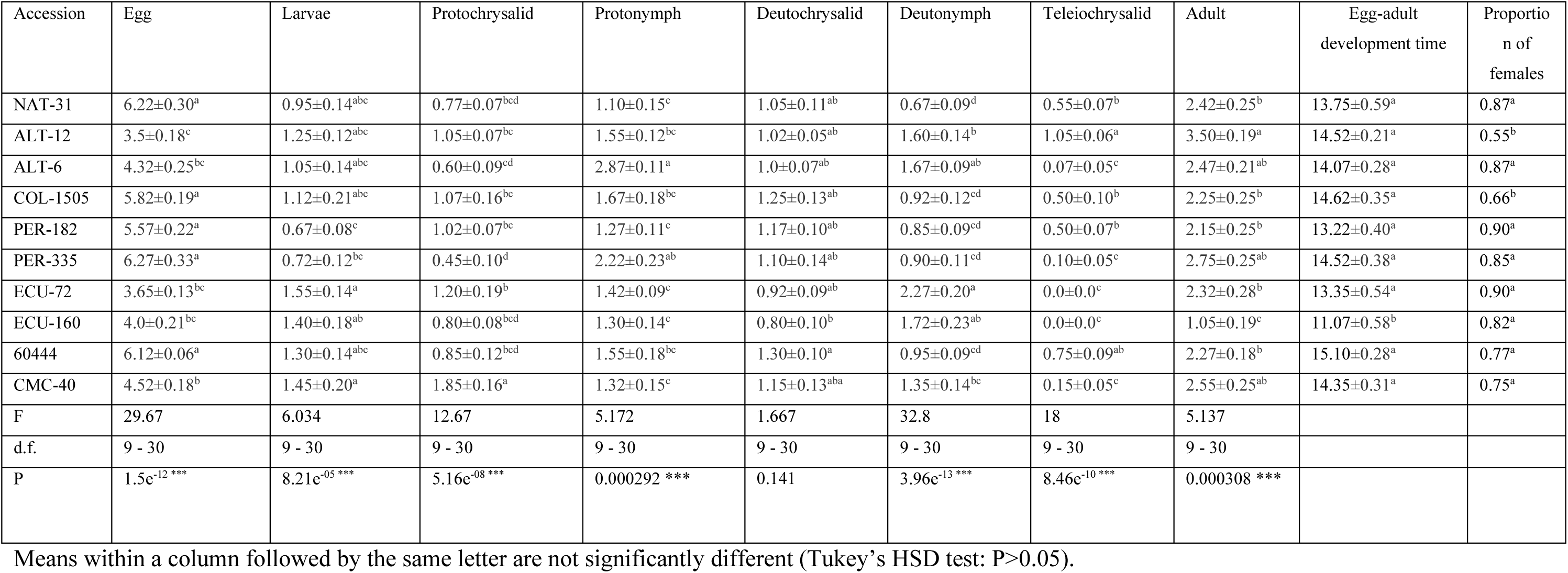
Time development per stages and proportion of *Mononychellus tanajoa* on selected cassava genotypes (*n*=50).

| Accession | Egg | Larvae | Protochrysalid | Protonymph | Deutochrysalid | Deutonymph | Teleiochrysalid | Adult | Egg-adult development time | Proportion of females |
| --- | --- | --- | --- | --- | --- | --- | --- | --- | --- | --- |
| NAT-31 | 6.22±0.30<sup>a</sup> | 0.95±0.14<sup>abc</sup> | 0.77±0.07<sup>bcd</sup> | 1.10±0.15<sup>c</sup> | 1.05±0.11<sup>ab</sup> | 0.67±0.09<sup>d</sup> | 0.55±0.07<sup>b</sup> | 2.42±0.25<sup>b</sup> | 13.75±0.59<sup>a</sup> | 0.87<sup>a</sup> |
| ALT-12 | 3.5±0.18<sup>c</sup> | 1.25±0.12<sup>abc</sup> | 1.05±0.07<sup>bc</sup> | 1.55±0.12<sup>bc</sup> | 1.02±0.05<sup>ab</sup> | 1.60±0.14<sup>b</sup> | 1.05±0.06<sup>a</sup> | 3.50±0.19<sup>a</sup> | 14.52±0.21<sup>a</sup> | 0.55<sup>b</sup> |
| ALT-6 | 4.32±0.25<sup>bc</sup> | 1.05±0.14<sup>abc</sup> | 0.60±0.09<sup>cd</sup> | 2.87±0.11<sup>a</sup> | 1.0±0.07<sup>ab</sup> | 1.67±0.09<sup>ab</sup> | 0.07±0.05<sup>c</sup> | 2.47±0.21<sup>ab</sup> | 14.07±0.28<sup>a</sup> | 0.87<sup>a</sup> |
| COL-1505 | 5.82±0.19<sup>a</sup> | 1.12±0.21<sup>abc</sup> | 1.07±0.16<sup>bc</sup> | 1.67±0.18<sup>bc</sup> | 1.25±0.13<sup>ab</sup> | 0.92±0.12<sup>cd</sup> | 0.50±0.10<sup>b</sup> | 2.25±0.25<sup>b</sup> | 14.62±0.35<sup>a</sup> | 0.66<sup>b</sup> |
| PER-182 | 5.57±0.22<sup>a</sup> | 0.67±0.08<sup>c</sup> | 1.02±0.07<sup>bc</sup> | 1.27±0.11<sup>c</sup> | 1.17±0.10<sup>ab</sup> | 0.85±0.09<sup>cd</sup> | 0.50±0.07<sup>b</sup> | 2.15±0.25<sup>b</sup> | 13.22±0.40<sup>a</sup> | 0.90<sup>a</sup> |
| PER-335 | 6.27±0.33<sup>a</sup> | 0.72±0.12<sup>bc</sup> | 0.45±0.10<sup>d</sup> | 2.22±0.23<sup>ab</sup> | 1.10±0.14<sup>ab</sup> | 0.90±0.11<sup>cd</sup> | 0.10±0.05<sup>c</sup> | 2.75±0.25<sup>ab</sup> | 14.52±0.38<sup>a</sup> | 0.85<sup>a</sup> |
| ECU-72 | 3.65±0.13<sup>bc</sup> | 1.55±0.14<sup>a</sup> | 1.20±0.19<sup>b</sup> | 1.42±0.09<sup>c</sup> | 0.92±0.09<sup>ab</sup> | 2.27±0.20<sup>a</sup> | 0.0±0.0<sup>c</sup> | 2.32±0.28<sup>b</sup> | 13.35±0.54<sup>a</sup> | 0.90<sup>a</sup> |
| ECU-160 | 4.0±0.21<sup>bc</sup> | 1.40±0.18<sup>ab</sup> | 0.80±0.08<sup>bcd</sup> | 1.30±0.14<sup>c</sup> | 0.80±0.10<sup>b</sup> | 1.72±0.23<sup>ab</sup> | 0.0±0.0<sup>c</sup> | 1.05±0.19<sup>c</sup> | 11.07±0.58<sup>b</sup> | 0.82<sup>a</sup> |
| 60444 | 6.12±0.06<sup>a</sup> | 1.30±0.14<sup>abc</sup> | 0.85±0.12<sup>bcd</sup> | 1.55±0.18<sup>bc</sup> | 1.30±0.10<sup>a</sup> | 0.95±0.09<sup>cd</sup> | 0.75±0.09<sup>ab</sup> | 2.27±0.18<sup>b</sup> | 15.10±0.28<sup>a</sup> | 0.77<sup>a</sup> |
| CMC-40 | 4.52±0.18<sup>b</sup> | 1.45±0.20<sup>a</sup> | 1.85±0.16<sup>a</sup> | 1.32±0.15<sup>c</sup> | 1.15±0.13<sup>aba</sup> | 1.35±0.14<sup>bc</sup> | 0.15±0.05<sup>c</sup> | 2.55±0.25<sup>ab</sup> | 14.35±0.31<sup>a</sup> | 0.75<sup>a</sup> |
| F | 29.67 | 6.034 | 12.67 | 5.172 | 1.667 | 32.8 | 18 | 5.137 |  |  |
| d.f. | 9 - 30 | 9 - 30 | 9 - 30 | 9 - 30 | 9 - 30 | 9 - 30 | 9 - 30 | 9 - 30 |  |  |
| P | 1.5e<sup>-12</sup> *** | 8.21e<sup>-05</sup> *** | 5.16e<sup>-08</sup> *** | 0.000292 *** | 0.141 | 3.96e<sup>-13</sup> *** | 8.46e<sup>-10</sup> *** | 0.000308 *** |  |  |

Means within a column followed by the same letter are not significantly different (Tukey's HSD test: $P>0.05$ ).

## DISCUSSION

Overall, our results showed that *M. tanajoa* had higher preference for genotype 60444 than CMC-40 (Tables 2 and 3, Figures 2 and 3). CMC-40 has been previously identified as the most susceptible host to arthropod cassava pests (Bellotti, 2002; Burbano, 2007; Bohorquez, 2009). The free-choice bioassay proposed in the present study has been instrumental to compare genotype preference. CMC-40 appeared to have properties that render this genotype attractive for female colonization and oviposition. The genotype 60444, a model African variety used in virology and biotech studies (Bull *et al*., 2009; Anjanappa *et al*., 2016), displayed even higher preference to green mite as compared to CMC-40.

In both choice and no choice assays the permanence had a similar trend and shows contrasting preferences for all accessions. Oviposition preference of free choice assays (Table 2) allows us to rank genotypes in seven groups (from highest to lowest) according their suitability to host acari eggs. Thus: 60444; PER-335; ALT-12-CMC-40; COL-1505-ECU-72; PER-182-ECU-160; ALT-6 and NAT31. In contrast, only three groups are identified from no choice assays: 60444 and ECU-160; PER335-PER182-ALT12-COL1505-ECU72 and ALT6-NAT31-CMC40. Thus, in the no choice assay experiment, disregard of the genotype, it was possible to confirm the acari preference to oviposit on a given genotype. Our results indicate 60444 as the most preferred genotype while ALT-6 and NAT31 were the less preferred. These results allow us to establish that the chosen accessions for this study were actually a representative sample from the germplasm bank with a high resistance variability affecting green mite fitness. Further work is required in order to identify the chemical and physical barriers of the susceptible and resistance genotypes in the response of the green acari attack. Among the former, pre-formed or constitutive agents could influence the adaptation of the green mite to this genotype; among the latter, secondary metabolites that influence attraction or repellence towards the genotype might be involved in susceptibility (Schoonhoven *et al*., 2005; Dicke and Baldwin, 2010; Piesik *et al*., 2011). All of these represent important issues to be analysed.

Hence, *M. tanajoa* is likely to use diverse strategies for host plant selection than for oviposition. A main strategy is the recognition of host, depending on the physicochemical properties of the surface of the leaves. During this process, mites may stay longer in one genotype than another, without indicating a clear host selection for oviposition. Another possibility is that the mite chooses a genotype exclusively to feed on but another to lay its eggs. This, actually, might represent a protection strategy. NAT-31 and ALT-6 were the least preferred genotypes where the females remain, however the first was more preferred to oviposit than the second (Tables 2, 3).

Developmental times in our study were similar to those observed by Yaseen and Bennet (1977) and Yaninek *et al*. (1984). In those studies, the developmental time of *M. tanajoa* was inversely proportional to temperature. Additionally, significant differences were present among the cassava genotypes at different stages of development (Table 4). Our study demonstrates a long development time in green mite (11.07-15.10 days) in all the genotypes, being only variable in ECU-160 (Table 4), which was the only genotype showing a significant difference compared to the other genotypes. Despite that no statistically significant difference was detected in the total developmental time of the green acari on the tested genotypes, differences were found when acari stages were analysed suggesting that each genotype exerts an antibiotic mechanism on the immature stages. An analysis of this variability shows that NAT-31, ECU-72 and ALT-12 genotypes appear to influence, either decreasing or increasing, developmental times of three different acari stages when compared to the rest of genotypes (ALT16, PER182, PER335, COL1505 and CMC40). In these cases, only a development stage was the influent, while in ECU160 two stages did. Arias (1995) and Gómez (2004) already reported a high mortality of nymphs when they fed on the ECU-72 material, concluding that the mortality of nymphal instars along with the length of the acari’s life cycle are a clear mechanism of antibiosis. In our study, it was observed that ECU-72 increases the time in stages of development such as larvae and deutonymph, while NAT-31 increases the egg stage, which might indicative of the influence of these genotypes by antibiosis at the time of development of the green mite. Table 2 shows the tendencies of genotypes NAT-31 and ALT-6 towards low oviposition as compared to the genotype 60444 which experiences higher levels of infestation. Our results are in agreement with previous observations of the green mite on CMC-40 system developed by Mesa *et al*. (1987) being mite behavior on CMC-40 similar to the parameters of oviposition and developmental time. As in the preliminary trial at both, choice and no choice assays, NAT-31 was identified as the less preferred, while 60444 was the most preferred in terms of green mite oviposition. NAT-31 is the less preferred probably because the disruption of oviposition prevents continuity of the mite progeny (see also Bohórquez, 2009; Carabali *et al*., 2009). Bohórquez (2009) suggests that NAT-31 shows antibiotic and antixenotic characteristics against *A. socialis*. However, Vargas *et al*. (2002) were the first to report the benefits of NAT-31, a variant of cassava (*M. esculenta*) resistant to whitefly (*A. socialis*) at the Valle Cálido of the Alto Magdalena. It is possible that resistance is due to the lineage as the parents of that genotype are ECU-72 and BRA-12. ECU-72 has previsouly been shown to be resistant to whitefly (Bohórquez, 2009) and recently reported to have high levels of green mite resistance. Likely, resistance established in the NAT-31 genotype comes from one of its parents (ECU-72) whose genes confer resistance to whitefly. Although the heritability of the resistance is unknown, most likely it is governed by several genes (polygenic). Hence, one can speculate that the other parent (BRA-12) presents some level of resistance to arthropods conferring an additive effect to NAT-31. However, there is a need to evaluate the possible resistance of BRA-12 to attack of the green mite. Genotypes ECU-72, ECU-160, PER-182 and PER-335 showed similar trends for mite oviposition preference and oviposition rate. These results mirror previous studies, which suggest that these genotypes are key elements in developing resistance to whiteflies (Burbano *et al*. 2007; Bohórquez, 2009; Carabalí *et al*., 2010a; 2010b; 2013), of these, NAT-31, ECU-72, 60444 genotypes were previously prioritized for in-depth screening (Burbano *et al*., 2007; Carabalí *et al*. 2010a, b). A particular response exists depending on the type of pest; in the case of green mites, one can hypothesise the capacity of adaptation and the ample host range the pests could prompt their coevolution with cassava. All of this might push the mite to behave in a polyphagous manner. Nevertheless more studies are needed on the host adaptability to *Manihot* relatives to evaluate and confirm the possible polyphagous mite behavior.

In all the cassava genotypes, *M. tanajoa* females began to oviposit within the first 24 h. Nearly 90-100% of the eggs had been oviposited onto the selected genotypes by the fourth day, (Figures 2 and 3). Since this oviposition rate illustrates the preference of the green mite towards different cassava genotypes, it is plausible to consider that cassava accessions show similar patterns of infestation and defensive responses in the first 24 h. Nevertheless, the high rates of oviposition recorded among the other genotypes were 35% lower than that observed in 60444. *M. tanajoa* females were observed to have high values oviposition preference and oviposition rates on the latter accession (Tables 2, 3 and 4).

In our study, NAT-31 displayed short development times for proto- and deutonymph, key stages in the mite’s life cycle. ECU-72 and ECU-160 tended to suppress the quiescent stages; hence, variants NAT-31, ECU-72 and ECU-160, suppressing and displaying development times, could possess an antibiosis mechanism that might partially explain various levels of resistance. In all genotypes, except ALT12 and COL1505, the proportion of females (0.9:0.1) was not affected (Table 4), within the established observations on male presented a constant motion seeking female’s quiescence states to fertilize. Another feature of the male is to feed very little when compared with females. It is important to note that the females decide which eggs will be male, due in large part to their haplo-diploid condition (Yaninek *et al*., 1988). Overall, results of the biology and preference for oviposition demonstrate that genotype NAT-31 exhibits resistance to *M. tanajoa*. Further, the different levels of resistance observed in the evaluated genotypes also suggest, variable levels of antixenosis and antibiosis. These findings are due mainly to the development time, suggesting that factor as responsible for the differences established between CMC-40 and the other genotypes. According to Sabelis (1985) changes in development time is the most crucial factor for the growth of mite populations.

In conclusion, oviposition preference, development time and sex ratio of the green mite were parameters allowing the identification of different levels of resistance (antixenotic and antibiosis) in the cassava germplasm.

Firstly, 60444 was the accession with levels of susceptibility higher than CMC-40 previously used in pest-cassava studies as susceptible check. Hence is plausible to conclude, in general, that 60444 can be considered as the most susceptible genotype indicating better adaptation of the mite to this host. The mite possesses a number of characteristics to assist in this behaviour, such as its high mutation rate and its aggregated distribution in the fields via colonies, which greatly reduces genetic crosstalk among organisms and makes resistance dilution difficult (Mesa *et al*., 1987; Saito *et al*., 1983). This response needs to be examined at the gene and molecular levels to explain the plant’s behaviour. Secondly, NAT-31 showed low population levels of *M. tanajoa*, which might indicate resistance to the green mite. This result could be exploited in genetic improvement programs for assisted selection of resistance to one of the most significant pests in the Americas and Africa (CIAT, 2006). Identification and characterization of accessions highly resistant to green mites will be particularly instrumental to investigate the molecular determinants of the resistance against green mite in cassava. Recent large-scale omics studies in cassava have helped to identify proteins associated with improved traits in cassava (Owiti *et al*., 2011; Vanderschuren *et al*., 2014). Similar studies with accessions contrasting for resistance against green mite (*i*.*e*. 60444, CMC-40, ECU-72 and NAT-31) could lead to the identification of genes, transcripts and protein expression patterns associated with pest resistance.

## Acknowledgments

Authors thank to Dr. Herve Vanderschuren for his valuable comments and help in editing versions of the manuscript. Also to Adriana Alzate, Adriano Muñoz and Mario Salazar for their technical support. The funds provided by Colciencias and Univalle throughout grants 1106-521-28281: ‘Molecular characterization of resistance in cassava (*Manihot esculenta* Crantz) to attack the green mite (*Mononychellus tanojoa*): a proteomics approach’ and the PhD grant awarded by Colciencias to the student Jaime Marín 4942009. To the editor, who did important recommendations that improved the original version of the manuscript.

